# SVR-based Multimodal Active Subspace Analysis for the Brain using Neuroimaging Data

**DOI:** 10.1101/2022.07.28.501879

**Authors:** Ishaan Batta, Anees Abrol, Vince D. Calhoun, the Alzheimer’s Disease Neuroimaging Initiative

**Author notes:** Data used in preparation of this article were obtained from the Alzheimer’s Disease Neuroimaging Initiative (ADNI) database (adni.loni.usc.edu). As such, the investigators within the ADNI contributed to the design and implementation of ADNI and/or provided data but did not participate in analysis or writing of this report. A complete listing of ADNI investigators can be found at: http://adni.loni.usc.edu/wp-content/uploads/how_to_apply/ADNI_Acknowledgement_List.pdf.

## Abstract

Understanding the patterns of changes in brain function and structure due to various disorders and diseases is of utmost importance. There have been numerous efforts toward successful biomarker discovery for complex brain disorders by evaluating neuroimaging datasets with novel analytical frameworks. However, due to the multi-faceted nature of the disorders involving a wide and overlapping range of symptoms as well as complex changes in structural and functional brain networks, it is increasingly important to devise computational frameworks that can consider the underlying patterns of heterogeneous changes with specific target assessments, at the same time producing a summarizing output from the high-dimensional neuroimaging data. While various machine learning approaches focus on diagnostic prediction, many learning frameworks analyze important features at the level of brain regions involved in prediction using supervised methods. Unsupervised learning methods have also been utilized to break down the neuroimaging features into lower dimensional components. However, most learning frameworks either do not consider the target assessment information while extracting brain subspaces, or can extract only higher dimensional importance associations as an ordered list of involved features, making manual interpretation at the level of subspaces difficult. We present a novel multimodal active subspace learning framework to understand various subspaces within the brain that are associated with changes in particular biological and cognitive traits. For a given cognitive or biological trait, our framework performs a decomposition of the feature importances to extract robust multimodal subspaces that define the most significant change in the given trait. Through a rigorous cross-validation procedure on an Alzheimer’s disease (AD) dataset, we show that our framework can extract subspaces covering both functional and structural modalities, which are specific to a given clinical assessment (like memory and other cognitive skills) and also retain predictive performance in standard machine learning algorithms. We show that our framework not only uncovers AD-related brain regions (e.g., hippocampus, entorhinal cortex) in the associated brain subspaces, but also enables an automated identification of multiple underlying structural and functional sub-systems of the brain that collectively characterize changes in memory and cognitive skill proficiency related to brain disorders like AD.

## 1 INTRODUCTION

In recent decades, it has become increasingly vital to understand the working of the human brain. With the advent of better and newer neuroimaging technologies, many efforts have been launched to improve our understanding of the structure and function of the brain. With more complexity in the neuroimaging data, computational methods for understanding the underlying patterns have been employed in numerous neuroimaging studies throughout the years. While on the one hand, studies have focused on enhancing the diagnostic capabilities using various learning models on the neuroimaging datasets (Klöppel et al., 2012), many studies have also been focusing on delineating the inherent structural and functional properties of the brain at various levels, especially in relation to finding biomarkers associated with various mental disorders as well as cognitive traits (Woo et al., 2017). Towards both these goals, numerous studies utilizing structural and functional magnetic resonance imaging (MRI) data have been conducted with various methods in the domain of machine learning.

One of the major themes in methods exploring the internal working of the brain has been to understand it in terms of brain regions/networks and their interactions, which could be arrived at by manual or data-driven selection methods (Calhoun et al., 2021; Mwangi et al., 2014). Features created at regional levels from neuroimaging data have been constantly used in various machine learning techniques to assess the important brain areas and connections towards a given prediction task associated with brain disorders (Sui et al., 2020). With the overall goal of developing relevant biomarkers, many such association studies have tried to identify indicators for various brain disorders. Given the variations involved in the sub-categories of brain disorders in terms of their overlapping diagnostic symptoms, it is increasingly difficult to relate the underlying changes in the brain to specific types of disorders (María Mateos-Pérez et al., 2018; Nielsen et al., 2020). While the set of brain regions involved in such disorders can be manually inferred from the outputs of learning models, more automated methods are useful to figure out the subsets of regions involved in various sub-categories or stages of brain disorders.

Subspace learning methods have been used in many contexts, including dimensionality reduction for high-dimensional analysis settings, enhancing discriminatory power for classification, transfer learning across datasets, subspace clustering for population analysis, and much more. Especially in image recognition tasks, subspace learning has been instrumental in handling high-volume and high-dimension data for classification purposes (Turan et al., 2021; Wang et al., 2020). Subspace learning frameworks have also been developed and employed for transfer learning by trying to preserve the inherent discriminatory subspace information across multi-corpus paradigms, including image (Razzaghi et al., 2019) as well as speech recognition tasks (Zhang and Song, 2019; Liu et al., 2018). Various deep subspace learning approaches have also been used in the context of classification (Sun et al., 2018) as well as subspace clustering (Abavisani and Patel, 2018; Lv et al., 2021).

However, in most cases, subspace learning approaches have been aimed toward generating low-dimensional features for either enhancing classification performance or clustering of datasets. Another important potential of using subspace learning is the fact that the learned low-dimensional representations of the data can be interpreted to represent certain meaningful components that define the feature set. In the context of the neuroimaging data, this would amount to the identification of various regional components of the brain from high-dimensional voxel-level data.

Towards this goal, many unsupervised techniques like independent component analysis (ICA) (Calhoun et al., 2009; Erhardt et al., 2011; Du et al., 2020) and principal component analysis (PCA) (Mwangi et al., 2014) have been used to extract meaningful data-driven subspaces from high dimensional neuroimaging data. While decomposing the data, they have been successfully used to reveal the underlying subspace patterns in the brain (Adali et al., 2018; Du et al., 2020). However, being unsupervised approaches, PCA- and ICA-based methods are not designed to take into account the diagnostic information and any related cognitive or biological assessment scores. Thus, the components generated with these methods are more general, and while they can be used for further association studies, they are not optimized to identify or encode information about brain disorders.

More recently, data-driven frameworks summarizing the associated biomarkers on a higher level in terms of brain-networks and brain-subdomains have enhanced the understanding of the working of the brain. Many studies have shown that the brain processes involve a well-connected set of networks even in the resting state, instead of individual regions (Fornito et al., 2015). In fact, resting-state studies have shown that functional connectivity within and between brain networks gets altered in various brain disorders (Fornito et al., 2015; Braun et al., 2018). Towards this goal, the use of multimodal neuroimaging data has also seen prominence in creating models that couple both structural and functional properties for studying and predicting various brain disorders (Calhoun and Sui, 2016). The changes in clinical indicators for brain disorders are often marked by underlying reorganization of the structural and functional properties of the brain. Determining the directions of such alterations in the brain in association with changes in cognitive or biological factors is key to defining successful biomarkers for brain disorders. Moreover, in most cases, there is a set of multiple brain sub-networks that undergo changes with the onset of various disorders, instead of a single sub-network or a single set of regions. This further calls for looking at these changes from the perspective of multiple subspaces instead of a single subset of features. Thus, it is essential to develop data-driven semi-supervised approaches that can reveal the brain’s underlying subspaces in terms of important trajectories of structural and functional changes that occur in association with given cognitive and biological traits.

Towards this direction, this work presents a new framework based on active subspace learning (ASL) (Ma et al., 2020; Constantine et al., 2014) to extract subspaces in the brain that co-vary together in association with a given trait at hand. We define the framework such that it can utilize multimodal information from structural and functional MRI data, resulting in subspaces that span both structural and functional aspects of changes in the brain. The framework is based on determining the significant directions of change from the gradient cloud of a regression function learned on a given dataset. This is achieved by the eigendecomposition of the covariance of the gradients of a learned mapping (regression function) from the input space of neuroimaging features to the output space of a target variable involving cognitive and biological scores. The prominent eigenvectors with significantly large eigenvalues correspond to the active subspaces capturing the essential directions of change in the brain features with respect to the target variable at hand.

We use support vector regression (SVR) as the underlying function for performing active subspace learning. While deep learning is also known to perform well for neuroimaging datasets (Abrol et al., 2021), in the context of meaningful active subspace learning, which provides better interpretability with lower-dimensional features, there are issues with deep learning models when using them as the underlying regression function. These are mainly due to deep learning models requiring high-dimensional voxel-level data as input instead of region-level measures for effective training, and the inability to establish direct correspondence between high-dimensional input features and the lower-dimensional representations learned in subsequent deeper layers. Thus, for the scope of this study, we use SVR, which is a standard machine learning regression method, as the underlying mapping function for the ASL analysis.

We also perform a repeated random sub-sampling procedure with held-out test data by aggregating and clustering active subspaces obtained from each repetition to make the analysis more robust. This clustering procedure estimates active subspace centers (ASCs) that essentially summarize important multimodal directions of change in the structural and functional properties associated with the target scores. We also show that when the input features are projected onto the active subspaces, the regression performance is retained, indicating that the active subspaces retain the predictive information while capturing the underlying subspace patterns in the brain. We run the analysis framework for the Alzheimer’s Disease Neuroimaging Initiative (ADNI) dataset with 7 cognitive and biological assessment scores to successfully analyze the underlying subspaces, collectively capturing the changes in the structural and functional components of the brain associated with various indicators for Alzheimer’s diseases (AD). Our results show that these subspace patterns are different when subspaces are computed for different groups within the dataset.

In essence, we show that our framework is successfully able to: (a) identify sparse and stable multimodal subspaces in the brain instead of determining only associated individual input features, (b) take into account the information from a given cognitive or biological trait when computing the subspaces rather than extracting generic unsupervised subspace patterns, (c) retain predictive information from the original input features while computing subspaces, and (d) provide with subspace signature patterns for the comparison of control subjects and subjects with specific brain conditions.

## 2 METHODOLOGY

### 2.1 Active Subspace Analysis

For a given point **u** ∈ ℝ^*m*^ in the space of the input data with *m* features, consider a function *f* : ℝ^*m*^ → ℝ that maps the input features from the input space to the space of the target variable *y*. The Active Subspace Learning (ASL) framework intends to compute important directions of covariance with the target variable in the form of collectively co-varying subspaces within the input space. Since the data in the input space by itself does not directly capture any direct information from the target variable, frameworks that work by decomposing the data points directly may not suffice for this aim. The ASL framework achieves this by performing decomposition in the gradient domain, i.e., the domain defined by the gradient ∇_**u**_ *f* of the function *f* with respect to **u**. Working in the gradient domain inherently takes into account the coupling of the input domain with the target domain while decomposing into subspaces. More specifically, this is achieved by the eigendecomposition of the expected covariance **C** of ∇_**u**_ *f*. **C** is defined as follows:

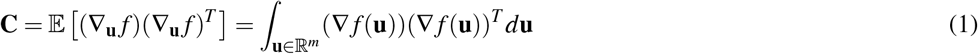

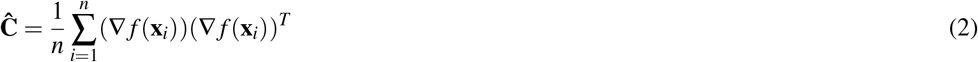

In most cases of learning-based approaches, **C** can be estimated as **Ĉ** from the data, which is effectively a sampling of the input space to use the gradients at each sampled point **x**_*i*_ ∈ ℝ^*m*^ for *i* ∈ {1, …, *n*} to calculate the estimator **Ĉ** as in Equation 2. In most cases, the defining parameters of the function *f* mapping the input and target variable space are learnt as part of the regression process from the dataset of sample size *n*, [**X**, **y**] with **X** ∈ ℝ^*m*×*n*^, **y** ∈ ℝ^*n*^. Once the function *f* is learned, the gradients and the expected covariance required to compute the matrix **Ĉ** can either be determined as a closed-form estimation or inferred from the data depending on the manner in which *f* is defined. Subsequently, the matrix **Ĉ** can be used as a close estimate for **C**.

It can be noted that the eigendecomposition of **C** would effectively capture the information of the average direction of the gradients. The eigendecomposition of **C** can therefore be used to define a set of active subspaces based on the eigenvectors corresponding to a significantly larger set of eigenvalues as below. These active subspaces, constituting the subset of “active” eigenvectors of **C**, can also be used to recreate a set of transformed features 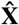 for further learning applications.

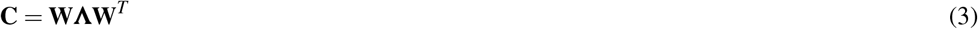

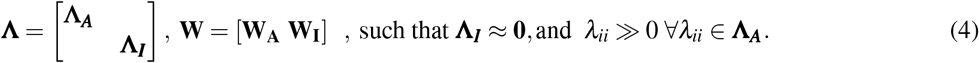

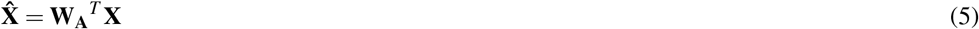

Essentially, the eigenvectors represented by the columns of **W** are the principal axes of variation in the gradient cloud of the function *f* based on the dataset [**X**, **y**], while eigenvalues in the diagonal matrix **Λ** capture the extent of this variation. Based on a threshold value of *λ*_0_ in the eigenvalue-spectrum, **W** is written as a horizontal concatenation of two matrices **W_A_** and **W_I_** representing active and inactive subspaces, respectively. The active subspaces signify the important directions in which the features in the input space co-vary together the most with variation in the target variable **y**. The directions with significantly higher importance are selected based on the corresponding eigenvalues to give active subspace vectors as in Equation 4.

Furthermore, the normalized eigenvalues can be used to define the fractional contribution of the corresponding subspace vectors towards the eigen-spectrum of subspaces. This can be used to measure the subspace vectors’ conducive strength toward the overall subspace structure. Using the vector form ***λ*** of the diagonal eigenvalue matrix **Λ** as ***λ*** = [Λ_11_, Λ_22_…Λ_*mm*_], the vector 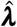 containing fractional eigenvalues is defined for the eigenvectors in the matrix **W** as follows:

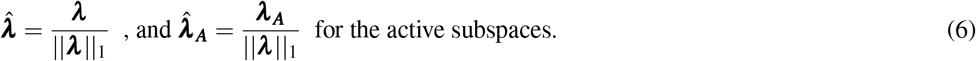

Note that the L1 norm here is the same as dividing by the sum of eigenvalues because the covariance matrix is symmetric and positive semi-definite by definition, resulting in all eigenvalues being positive.

### 2.2 Application to Neuroimaging Data

This analysis can have two-fold usage. Firstly, the transformed features 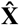 are then used for further learning to get potentially better prediction performance. Secondly, if a given active eigenvector represented by a column of **W_A_** is sparse, we can identify features (e.g., components or brain regions in structural magnetic resonance imaging (sMRI) data, functional connectivity in case of functional MRI data) that together form the active subspace corresponding to that eigenvector. This subspace can be said to be associated with the target variable **y** (which in this case is a specific cognitive or clinical score). In terms of structural MRI features (Xu et al., 2009), this can be interpreted as identifying structural subspaces (subsets of regions) co-varying the most with changes in a given cognitive trait. For static functional network connectivity (SFNC) features (Jafri et al., 2008), this can be interpreted as identifying functional subspaces (subsets of connections or sub-networks) in the brain whose change is associated with the prediction of the given score. For a combined multimodal input with both structural and functional features, one could obtain a multimodal subspace vector with both structural and functional elements, representing the collective changes from both modalities.

### 2.3 SVR-based Active Subspace Analysis

The gradient of the function *f* is one of the primary measures used to define the active subspace learning framework. In practical settings, it is not possible to determine the real function *f* underlying the relation between the input and target variables, but *f* is estimated by fitting a regression model to the dataset [**X**, **y**]. Previous methods have used Gaussian process regression (GPR) for applications in sensitivity analysis (Wycoff et al., 2021). However, these were applied to a relatively lower dimension (of up to 10) of data. For handling higher dimensional neuroimaging features, we employ kernel-based support vector regression (SVR) function as the mapping function for the aforementioned ASL analysis. The predictive model of SVR described as follows can be used as *f*,

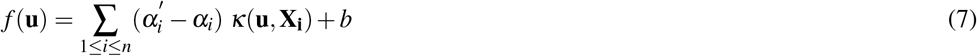

Where 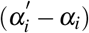 are the the Lagrange multipliers (dual coefficients) in SVR, *b* is the intercept term and *κ*(**u**, **X_i_**) is the kernel function used. The gradient of the learned predictive SVR model can thus be written as:

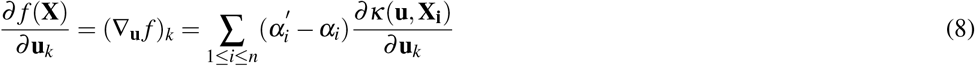

A widely used choice of kernels for the SVR function is the radial basis function (RBF) kernel because of its stationary, isotropic, and infinite smoothness properties. The RBF kernel is defined as 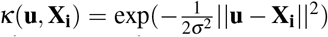. Previously, (Ma et al., 2020) employed SVR with RBF kernel for identifying subspaces for sensitivity analysis, but on relatively lower dimensional data with 16 variables. However, for higher dimensions, the exponential term involved in the RBF kernel expression causes numerical issues in the computation (Wang et al., 2018), resulting in negligible gradients, making it unsuitable for high-dimensional data involved in neuroimaging datasets. While the linear kernel is computationally feasible, it renders a constant gradient for every point, making it too simplistic for subspace analysis. The case of RBF and linear kernels is explained in further detail in the supplementary materials. We develop our framework using polynomial kernel-based SVR as the underlying function for computing active subspaces to avoid these possible numerical issues. For the polynomial kernel defined as 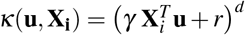, the gradient computation of the prediction function *f* is done as follows:

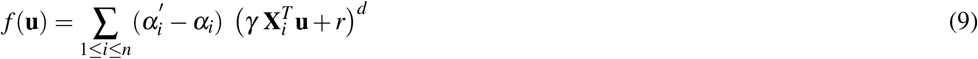

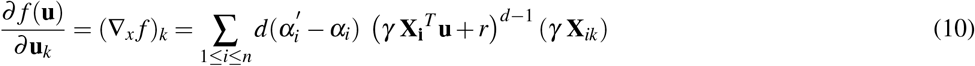

It should be noted that while the gradient expression in the case of the polynomial kernel in Equation 10 is theoretically derived from the SVR regression function, the regression function is itself learned from the data in terms of the involved Lagrange multipliers 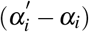 and the intercept term *b*. Thus, it mixes a theoretically derived gradient expression upon a function learned from data. These gradient value estimates at various data-points can be used to compute the sample covariance matrix **Ĉ** in Equation 2, resulting in an estimate of matrix **C** in Equation 1.

### 2.4 Repeated Sub-sampling to Extract Robust Active Subspace Centers

The active subspace analysis done on multiple random test-train splits of the data can be used to create a set of active subspace matrices 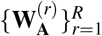 and their corresponding fraction contributions to the subspace eigen-spectrum 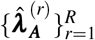 as defined in Equation 4 and Equation 6 respectively, each consisting of only the active subspace vectors and their fractional contributions from a given repetition (*r*) of the analysis. These vectors, taken together across repetitions, can be clustered using a standard clustering algorithm such as k-means, i.e., clustering is performed on the set of vectors obtained by aggregating active subspace vectors from all the *R* repetitions into one matrix 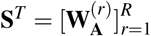. The matrix 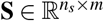 collects the set of *n_s_* eigenvectors (*n_s_* ≥ *R*), which summarize the spectral properties of ∇_**u**_ *f* via repeated sub-sampling of the training dataset. Correspondingly, the vector 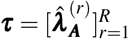 captures the fractional importance of the aggregated eigenvectors in **S** in their respective eigen-spectra.

Since performing multiple eigendecompositions is known to be affected by sign ambiguities (Bro et al., 2008), alignment of each vector in **S** was done with the gradient cloud for its corresponding repetition such that its dot product with the majority of the gradient points is positive, following a procedure similar to (Bro et al., 2008). K-means clustering is then performed with each row of **S** as a data-point, optimized for the best number of clusters *n_c_* using the silhouette coefficient metric. To remove any further possible sign ambiguities, the correlation between cluster centroids was computed, and pairs with a high anti-correlation (> 0.9) were considered as duplicates due to sign ambiguity. From the obtained cluster labels 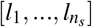 for each eigenvector in **S**, the centroids 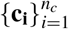 were computed for all the clusters as:

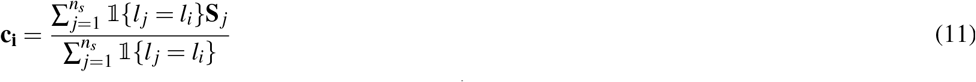

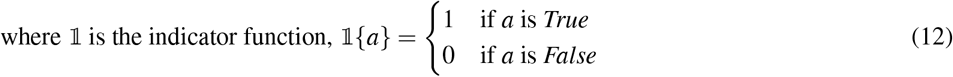

The L2-normalized active subspace centers defined by the cluster centroids for the most discrete clusters essentially summarize across repetitions the principal directions in which the features from the data co-vary with the target variable, as represented by the active subspace vectors.

Finally, for ranking the top active subspace centers, the mean fractional eigenvalue, as the mean contribution of their constituent eigenvectors to their corresponding eigen-spectra, was used as the measure ascribing importance. For each of the *n_c_* clusters, the mean fractional contribution 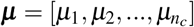 vector for the *n_c_* clusters is defined as:

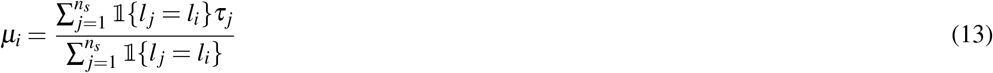

Thus, using the repeated sub-sampling procedure for the extraction of active subspace centers in the form of cluster centroids 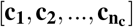 and the corresponding mean fractional contributions of active subspace centers (**μ**), one can determine and rank the important directions in which structural and functional measures from the brain co-vary collectively in association with scores from a given trait.

## 3 EXPERIMENTS AND RESULTS

### 3.1 Dataset and Preprocessing

Data used in the preparation of this article were obtained from the Alzheimer’s Disease Neuroimaging Initiative (ADNI) database (adni.loni.usc.edu). The ADNI was launched in 2003 as a public-private partnership, led by Principal Investigator Michael W. Weiner, MD. The primary goal of ADNI has been to test whether serial magnetic resonance imaging (MRI), positron emission tomography (PET), other biological markers, and clinical and neuropsychological assessment can be combined to measure the progression of mild cognitive impairment (MCI) and early Alzheimer’s disease (AD).

We used functional and structural MRI data from the Alzheimer’s Disease Neuroimaging Initiative (ADNI) dataset. The preprocessing was done as in (Abrol et al., 2020), and only first visit scans for subjects with available structural MRI, functional MRI, and clinical scores were used for the analysis. The fMRI data were preprocessed via an SPM12 pipeline involving rigid body motion correction to correct subject head motion, slice-timing correction, warping to the standard MNI space using the EPI template, resampling to (3mm)^3^ isotropic voxels, and smoothing using a Gaussian kernel (FWHM = 6mm). The sMRI data preprocessing included segmentation using modulated normalization algorithm, following which we smoothed the gray matter volume (GMV) maps using a Gaussian kernel (FWHM = 6 mm). Quality control (QC) of the preprocessed sMRI and fMRI datasets included discarding images that exhibited low correlation with individual and/or group level masks. Additionally, for fMRI data, images with high head motion were discarded to rule out potential spurious differences in functional connectivity. The seven clinical scores used include age, Mini-Mental State Exam (MMSE), Functional Activities Questionnaire (FAQ), Clinical Dementia Rating Scale Sum of Boxes (CDRSB), Alzheimer’s Disease Assessment Scale (ADAS11, ADAS13), and Rey’s Auditory Verbal Learning Test (RAVLT). After selecting subjects with both sMRI and fMRI data as well as all the scores available, a total of 606 subjects with 3 groups were used. These groups included 217 control subjects (CN), 324 subjects with mild cognitive impairment (MCI), and 65 subjects with Alzheimer’s disease diagnosis (AD). More information about the demographics of the data is included in the supplementary materials.

### 3.2 scICA Component Maps

From a fully automated spatially constrained ICA (scICA) framework on fMRI data based on the Neuro-mark study (Du et al., 2020), 53 spatially independent components with a high correlation threshold for multi-dataset correspondence were obtained (Du et al., 2020). The pre-computed Neuromark components available in the GIFT toolbox (https://trendscenter.org/software/gift/) were used. Table 1 shows the details with the names and MNI coordinates for brain regions in these component maps. The fMRI time courses of the 53 components estimated from the scICA framework were used to compute the functional connectivity features. More specifically, static functional connectivity (SFNC) was estimated as pairwise Pearson’s correlation coefficient of these 53 component time-courses, leading to a total of 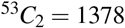 SFNC features for each subject. Likewise, structural masks of the components were mapped to the structural domain to extract the 53 structural GMV features for each subject, with each feature representing the mean value across component voxels in the brain mask.

**Table 1.**
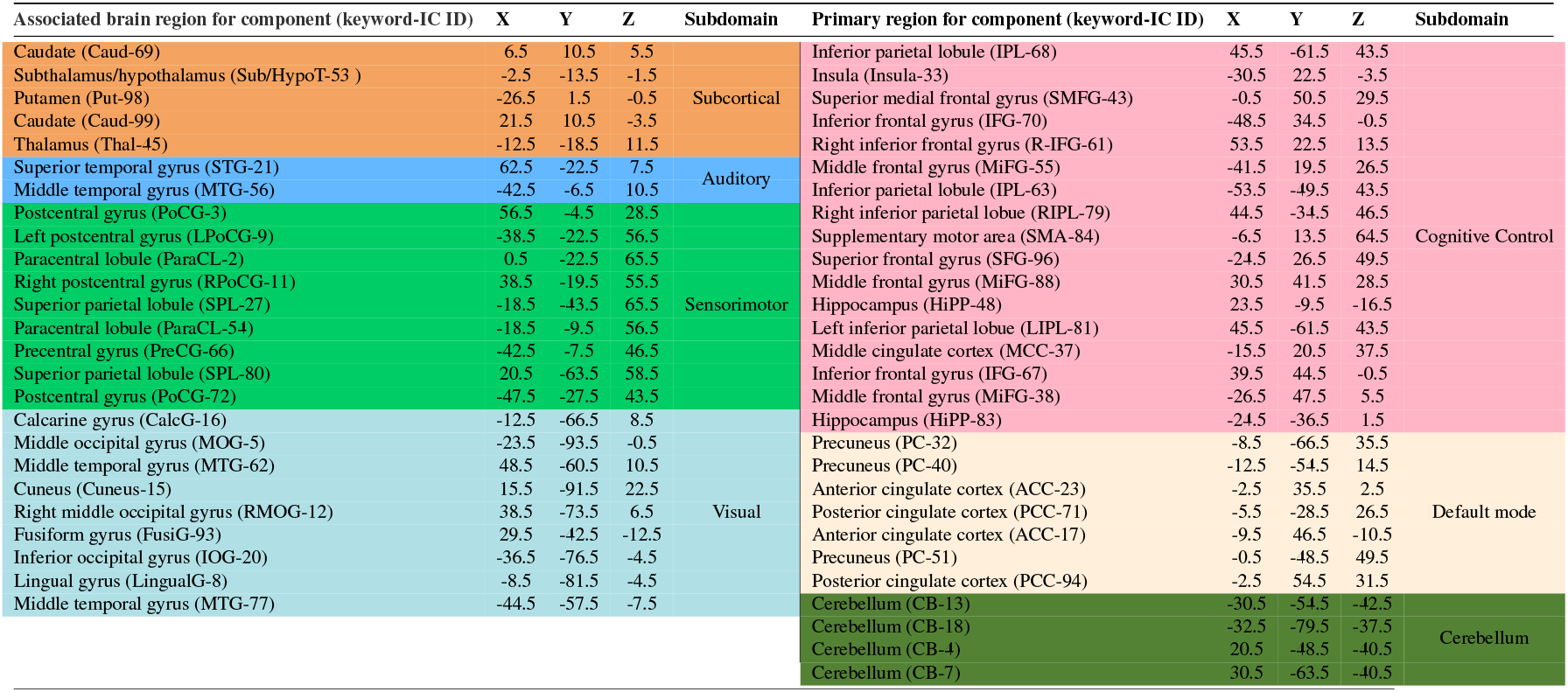
Table showing the peak MNI coordinates and the associated brain regions for the 53 scICA components extracted in the Neuromark framework (Du et al., 2020) on the fMRI data. The 53 components belonging to 7 subdomains (resting-state networks) are listed in distinct colors in the table, and are available in the GIFT toolbox (https://trendscenter.org/software/gift/) under *icatb_templates/neuromark_53*. The keyword for each component is shown in the parenthesis against the component name, and is used for addressing them in subsequent figures in this paper. These components were further used to create structural masks in the MNI domain which were used to create 53 structural features from the GMV maps of the subjects. The functional connectivity features along with the structural features were used as input to the active subspace framework (See Figure 1).

**Figure 1.**
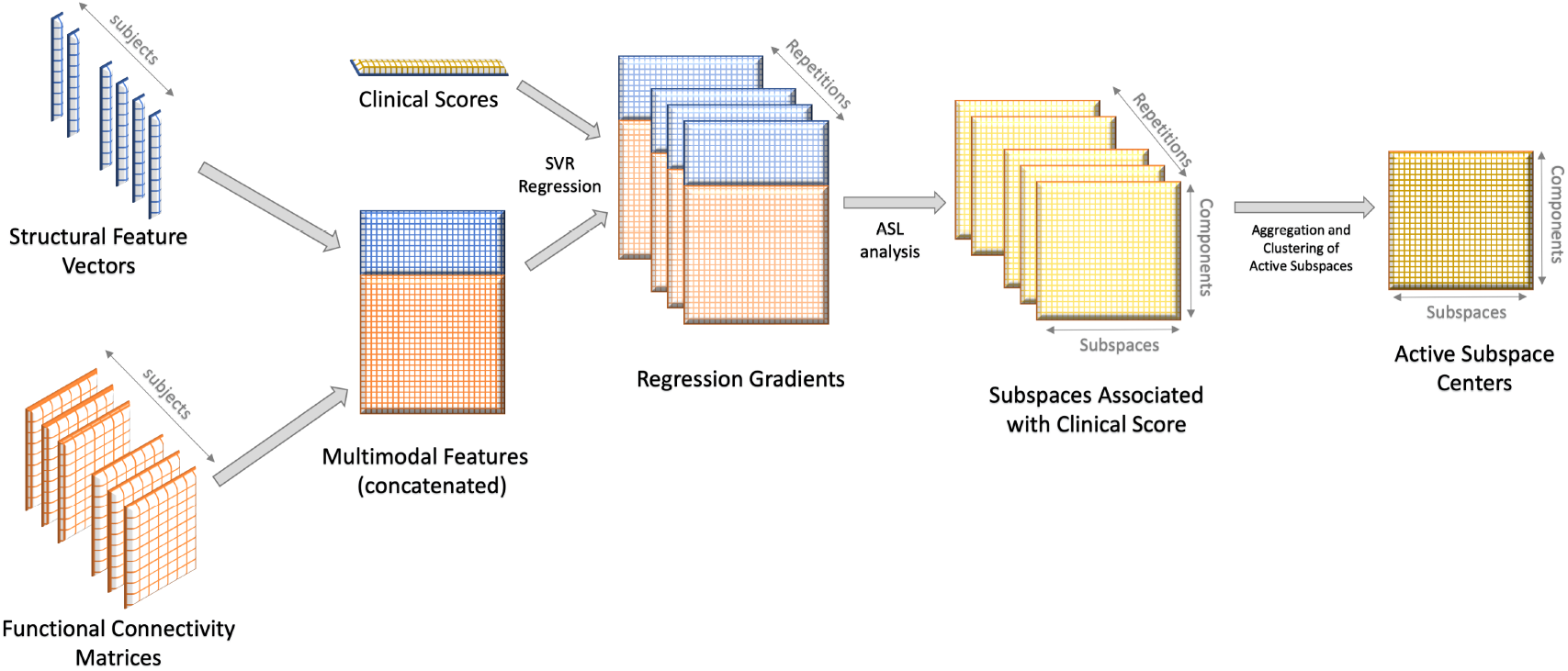
A schematic diagram showing the step-by-step procedures involved in the overall approach to extract Active Subspace Centers (ASCs) using the Active Subspace Learning (ASL) framework. Starting with fused structural and functional features from brain components as input, ASL is performed for 100 repetitions with support vector regression as the underlying function, followed by the aggregation and clustering of active subspace vectors across repetitions to extract the ASCs.

**Figure 2.**
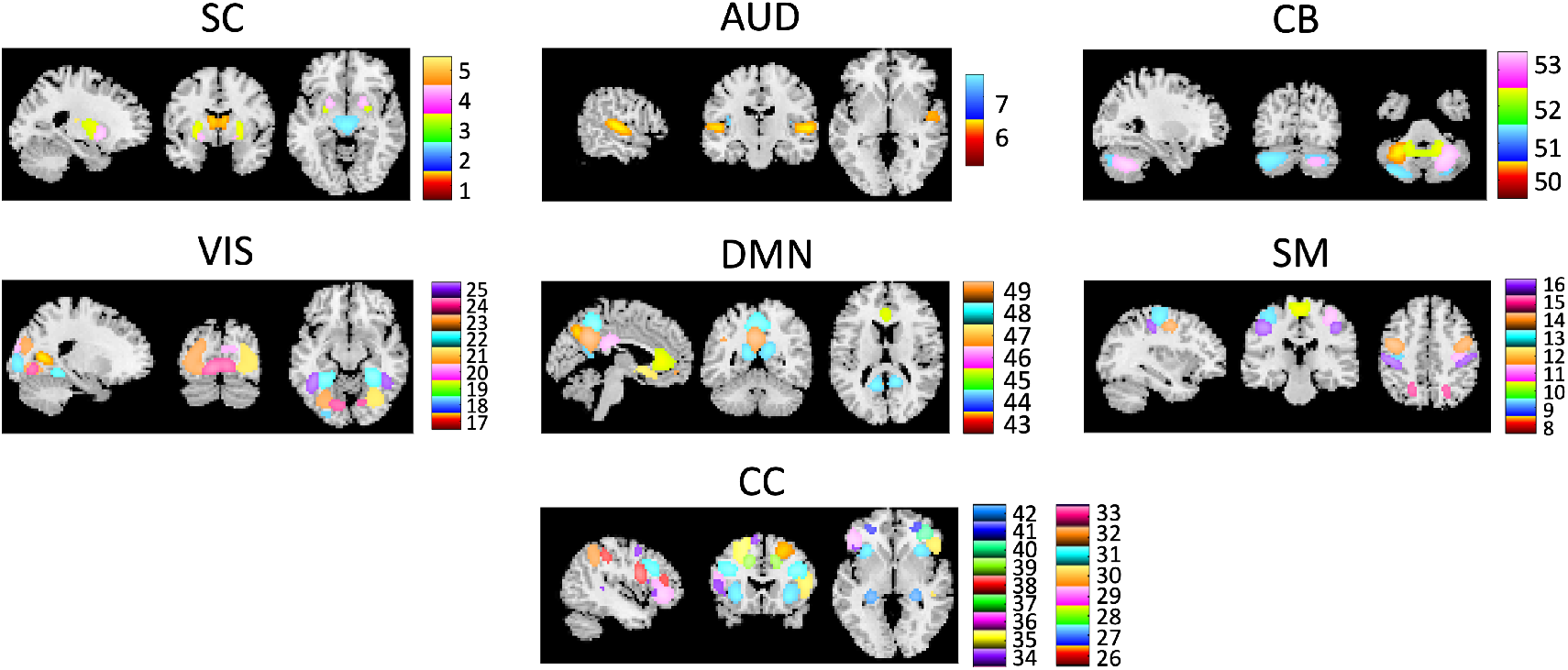
A total of 53 components corresponding to various brain regions extracted by the scICA procedure using the Neuromark framework are shown in the figure. The components are shown after dividing into 7 brain subdomains (resting state networks), namely Subcortical (SC), Auditory (AUD), Sensorimotor (SM), Visual (VIS), Cognitive Control (CC), Default Mode Network (DMN), and Cerebellar (CB) areas.

### 3.3 Existence of Active Subspaces

The SVR-based multimodal active subspace learning framework described in subsection 2.1,2.3 was used on the joint structural (GMV), and functional (SFNC) features. As mentioned in subsection 2.4, the framework was run for 100 repetitions, each with randomly selected 80% of the subjects. For each repetition, firstly, an SVR model with a polynomial kernel was trained with 5-fold internal cross-validation on 80% of the randomly selected training set. For computation and comparison of subspaces based on various groups of subjects in the data, the whole analysis was performed separately with subjects only from each of the groups (CN, MCI, AD) as well as with the full dataset (global group). Since the dimensions of the structural and functional features are 53 and 1378 respectively, we selected the top 100 SFNC features based on the weights of training a unimodal linear SVR model on the SFNC data, resulting in 153 total features per subject comprising of 53 structural GMV and 100 SFNC features. These features were used jointly as input to the ASL model leading us to compute an active subspace matrix 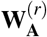 for each repetition *r* = 1, 2, .., 100.

Subsequently, the set of active subspace matrices from all repetitions was used to create the matrix **S** consisting of active subspace vectors from the columns all matrices in 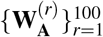. The matrix **S** was constructed to inspect the existence of and also compute any consistently featuring directions in the eigen-spectra of the gradient clouds from the repeated analysis. Towards this, k-means clustering of eigenvectors (represented by columns of the matrix **S**) was done to check for these consistent directions. Since there are known sign ambiguity issues with eigendecomposition (Bro et al., 2013), the signs of the vectors in the matrix **S** were aligned towards the sign of the majority of the points in the gradient cloud before performing the clustering step. This approach has been previously used by (Bro et al., 2008) for singular value decomposition and ensures in our case that no two clusters obtained are representing the same principle direction of the gradient cloud but with opposite signs. Thus, if discrete clusters are obtained upon subsequent clustering, they could be considered as signifying a consistent set of directions in the form of active subspaces.

The t-Distributed Stochastic Neighbor Embedding (tSNE) plots for the clustering of active subspace vectors in **S** are shown in Figure 3 for analysis on subjects exclusively from each grouping (global, CN, MCI, AD). The clustering model order was optimized based on the average silhouette coefficient for *k* = 2, 3, .., 50 for the k-means algorithm. For all the 28 cases of 4 groupings and 7 scores, the mean silhouette coefficient was higher than 0.1 for the optimized model order. As is also clear from the multiple discrete clusters obtained in Figure 3, the subspace structure in all the 28 cases does contain several active subspaces consistently featuring in the repeated analysis on randomly selected training subsets of the data.

**Figure 3.**
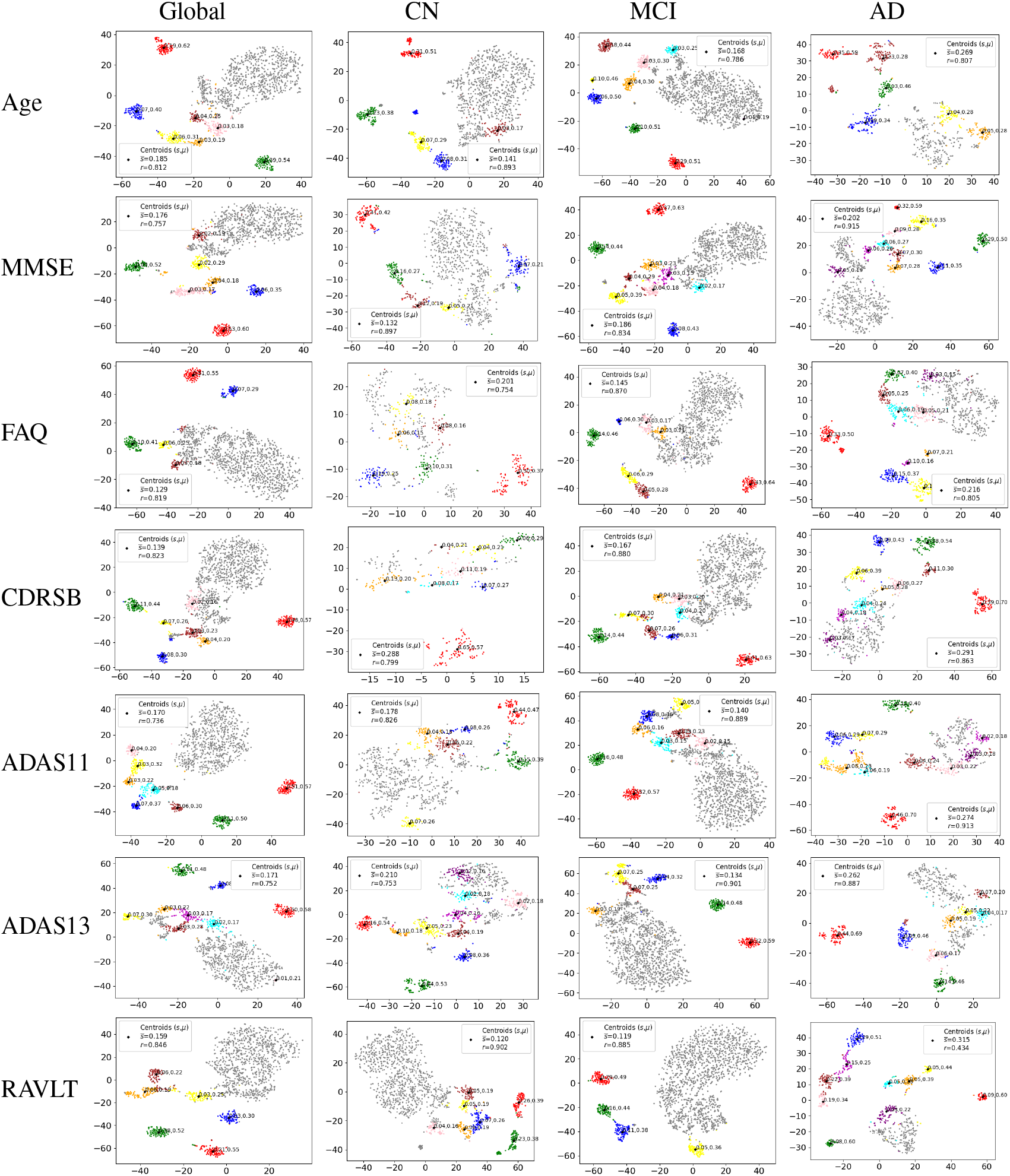
t-SNE embeddings are visualized for the active subspace vectors aggregated from across 100 repetitions of the ASL analysis. K-means clustering was performed on the aggregated vectors for each of the cases of the data from the seven scores and four groupings of the dataset (28 total cases). Top clusters (shown in different random colors) in each case were selected based on the mean fractional contribution (*μ* > 0.1) of the constituent vectors in that cluster (See subsection 2.4). As observable in all cases, the existence of discrete clusters reveals the existence of important directions which are consistently present across multiple repetitions of the analysis on randomly selected subsets of the data. The centroid for each of the discrete clusters is a multimodal vector taken as the active subspace center (ASC), encoding an important direction of collective changes in brain components and connections, which are associated with changes in the target score. (*μ*: mean fractional contribution, *s*: mean silhouette score for an individual cluster; 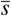: average silhouette score across all clusters, *r*: correlation between *μ* and *s* values of the clusters. Since the correlation (*r*) values are high in each case, using *μ* as a metric does not compromise on cluster separation captured by *s*.)

### 3.4 Active Subspace Centers

As described in subsection 2.4, the mean *μ_i_* of the fractional contribution of the subspace eigenvectors belonging to each of the clusters to the eigen-spectrum of their corresponding repetition was used to rank the cluster centroids 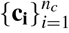 representing multimodal active subspace centers (ASCs). Essentially, *μ* is a measure to capture whether the subspace centers comprise similar active subspaces from various repetitions. Figure 3 shows all the clusters and their centroids (active subspace centers) for the ASL analysis on each of the 4 groups (global, CN, MCI, AD) with each of the 7 scores as the target variable. Based on the value of *μ_i_* of the *i^th^* subspace center, the subspace centers with *μ_i_* > 0.1 were considered as active subspace centers, representing a significant contribution of the constituent eigenvectors of the clusters to their corresponding eigen-spectrum. The threshold of *μ* > 0.1 for a cluster centroid (subspace center) to be considered as an active subspace center was selected so that the mean fractional contribution value is sufficient to be contributing to the 90^*th*^ percentile of the eigen-spectrum. It can be noticed that for each case, the total number of optimized clusters obtained may be different based on the silhouette score-based optimization procedure used for finding the number of optimal clusters for the k-means clustering procedure.

Figure 4 shows all the ASCs that were selected based on a value of *μ* > 0.1 for the global group. A similar plot for other groups (HC, MCI, AD) can be found in the. Figure 5 shows a group-wise layout for the highest ranked active subspace center (ASC) for all the cases of 4 groups and 7 scores. Since the ASCs are multimodal vectors with both structural as well functional features, they are visualized as a multimodal connectogram with the nodes representing the structural components and edges representing the functional connection features between them. The color of the nodes and edges represents the values from the ASCs for the corresponding structural and functional features, respectively. It should be noted that the ASCs are unit norm multimodal direction vectors representing the direction in which the multimodal brain features show the highest change in association with the target variable at hand. As is clear from Figure 5, our framework is able to extract multimodal subspaces that are sparse and expressed in terms of subsets of combined structural and functional features. Notably, there are significant differences in multimodal features that constitute the most active subspace centers for various groups for each of the given scores, indicating that the underlying subspaces in the brain differ according to the various stages of Alzheimer’s disease.

**Figure 4.**
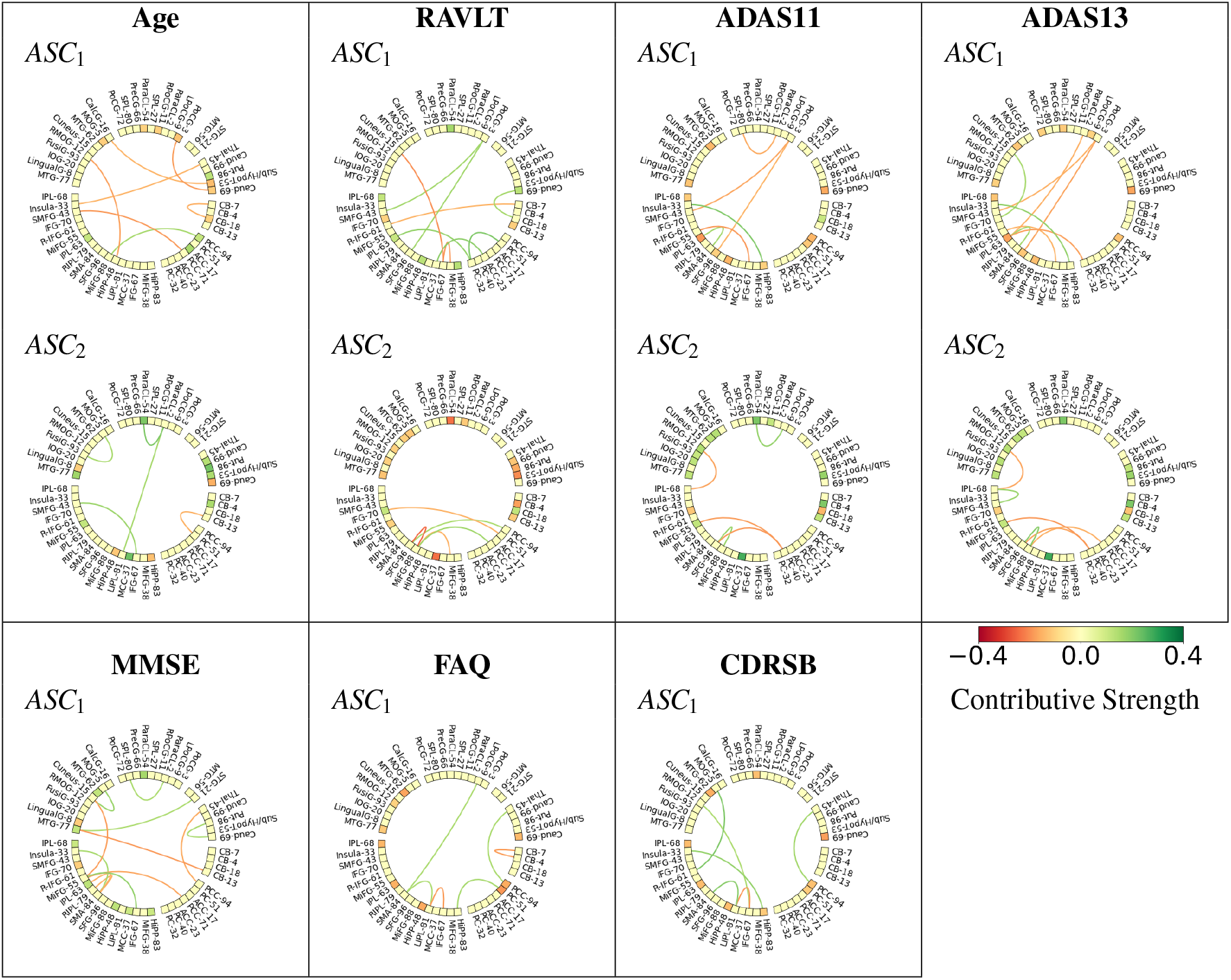
The most contributing multimodal active subspace centers (ASCs), ranked according to the mean fractional contribution (*μ*) with *μ* > 0.1, computed on all subjects, are shown for the seven scores. Each ASC is represented by a unit norm multimodal direction vector with elements visualized as a multimodal connectogram. The node and edge colors signify the values of the structural and functional elements of the multimodal ASC, respectively. Moreover, it can be noted that the contributive strength for memory-related regions like the hippocampus (Hipp-48, Hipp-83) is negative for scores known for higher values with a decline in memory performance (age, FAQ, CDRSB, ADAS), which is in turn related to shrinkage in these areas. Along similar lines, scores like RAVLT and MMSE, which have a higher value for a better performance, show a positive contribution from the hippocampus in the ASCs. This is in line with the expected contribution of the hippocampus, given its role in memory-related tasks and structural as well as functional changes involved with cognitive decline in Alzheimer’s disease.

**Figure 5.**
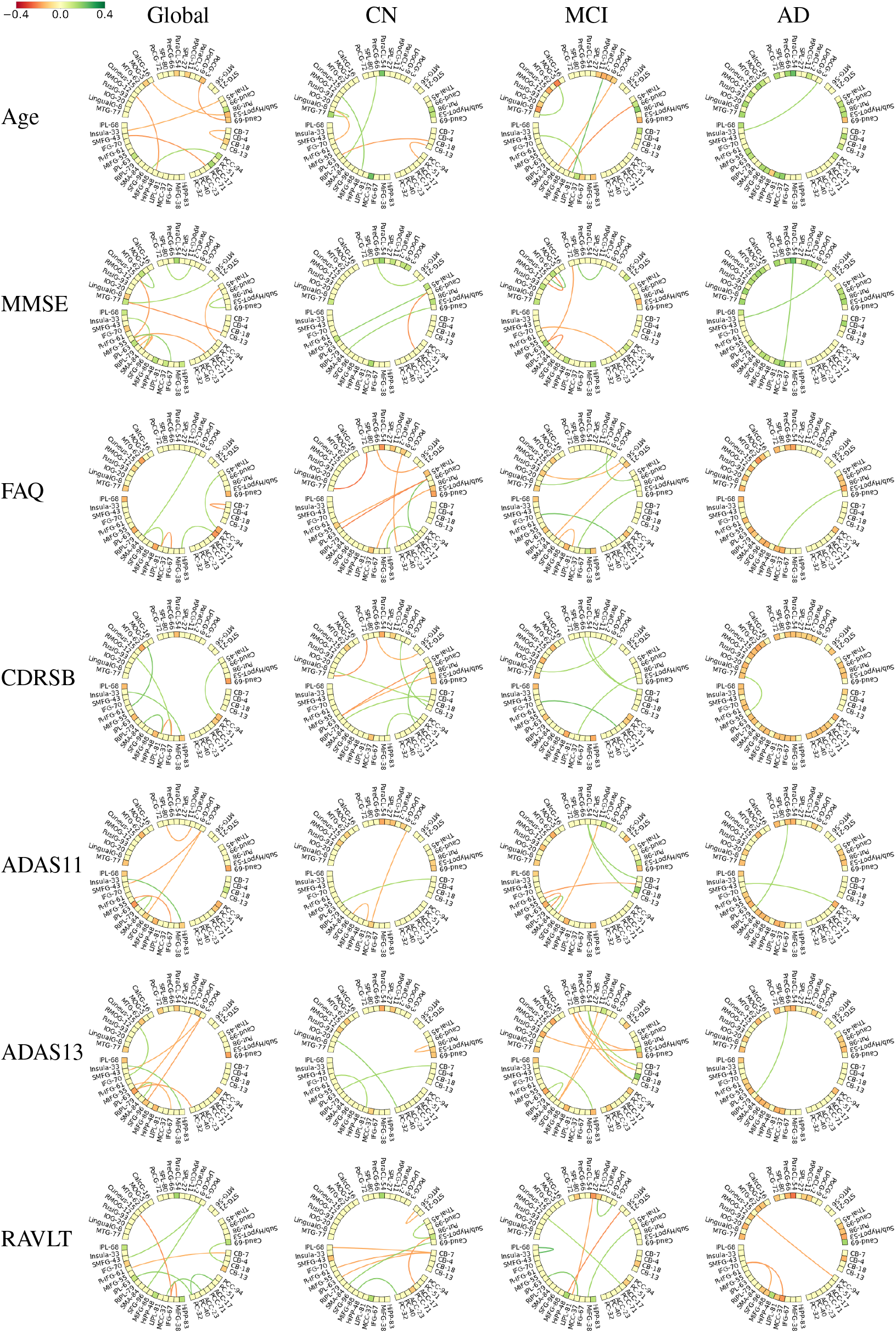
Top multimodal active subspace centers (ASCs). Each ASC is a unit norm multimodal direction vector with elements visualized as a multimodal connectogram with node and edge colors showing the values of contributive strength of the brain components and connections from the structural and functional parts of the vector. It can be noted that in line with the findings of previous MRI studies on Alzheimer’s disease having greater sensitivity for brain structure compared to function, structural features are the main contributors to the multimodal ASCs corresponding to the AD group, unlike the other groups.

### 3.5 Biological Findings

Figure 4 shows the top multimodal active subspace centers (ASCs) for each of the seven scores used in the analysis. Many previous studies have been done to find the association of individual brain regions and connections with ageing and Alzheimer’s disease. However, with the ASL framework, it is possible to decompose the association pattern to get underlying subspaces that collectively characterize their relationship with the target variables at hand. Based on the results depicted in Figure 4, it can be noticed that many of the regions that define the top ASCs are previously known to have associations with the target scores. For example, the structural and functional connectivity of the hippocampus is known to be affected in Alzheimer’s disease. This is also observable with respect to its negative contribution in the ASCs corresponding to age, FAQ, ADAS, and CDRSB, all of which are indicators of higher dementia, which in turn is related to shrinkage in the hippocampal volume and connectivity. Along similar lines, RAVLT and MMSE scores, which have a higher value for better performance in memory and impairment-related tasks, the contributive strength of hippocampus areas was positive (Figure 4). For the RAVLT score involving the performance for episodic memory recall, a positive contribution of the hippocampus components was observed, in line with its known role in being positively associated with episodic memory and RAVLT score prediction (Moradi et al., 2017).

Ageing is known to affect various brain systems, including memory and cognitive control. Regions that define the ASCs corresponding to changes in age were hippocampus, putamen, middle temporal, post-central, superior, and mid-frontal gyri, which are in line with previous findings about age-related changes in gray matter volume and functional connectivity (Salami et al., 2012; Nyberg et al., 2020). For the functional part of the multimodal ASCs, connections involving the hippocampus, thalamus, inferior frontal, superior frontal, insular, calcarine, and fusiform areas had major contributions.

The MMSE score (Folstein et al., 1975; Molloy and Standish, 1997), which is a composite measure of orientation, attention, memory, language, and visual-spatial skills, has been studied well for underlying associations of brain regions (Zhang et al., 2012; Bhagwat et al., 2019; Wan et al., 2012; He et al., 2007; Golbabaei et al., 2016). Changes in MMSE score are known to be associated with changes in structural and functional properties of various brain regions, including the hippocampus, middle-superior temporal (Zhang et al., 2012) and inferior frontal areas (Wan et al., 2012). All of these areas feature in the ASCs associated with the MMSE score. For the RAVLT score, regions featuring in the ASCs like middle/inferior occipital, middle cingulate, inferior occipital, inferior frontal (pars triangularis), inferior parietal, and middle temporal gyri (entorhinal cortex) are all known to be individually associated with RAVLT score prediction (Moradi et al., 2017; Wan et al., 2012; Gomar et al., 2011). Regions associated with the FAQ score included areas from the hippocampus, entorhinal cortex, inferior parietal areas, precuneus, and middle temporal gyrus. Many of these areas have previously known associations with FAQ scores and also with predicting the transition into AD during the MCI stage (Gomar et al., 2011).

The CDRSB score (Hughes et al., 1982) is measured based on a combination of various cognitive and behavioral domain functions like memory, orientation, judgment and problem solving, community affairs, home, and hobbies performance, and personal care. Different values of this score are also used to categorize patients into healthy, MCI, and AD conditions (Hughes et al., 1982). Regions in the ASCs associated with CDRSB involved areas from hippocampal, caudal, middle frontal, superior frontal, inferior parietal, and medial temporal gyri, as well as the posterior cingulate cortex (PCC). Previous studies on the association of CDRSB with neuroimaging features (Chang et al., 2011) have also found a similar set of areas, including frontal, temporal, parietal, and PCC regions (Perneczky et al., 2007) to be associated with thinning of the cortex in MCI groups.

ADAS rating (Rosen et al., 1984) consists of both cognitive and non-cognitive sections covering language, memory, praxis, and orientation. Hippocampus, middle temporal, entorhinal cortex, inferior parietal, caudate, and middle occipital gyrus areas were the primary contributors to the multimodal ASCs for ADAS scores. Many of these areas have previously known associations with changes in ADAS-cog score (Zhang et al., 2012; Gomar et al., 2011; Bhagwat et al., 2019). Interestingly, ASCs associated with ADAS11 and ADAS13 scores are highly overlapping in figures 4 and 5 in terms of the contributing brain components and connections. Since both these scores are very similar to each other, this further points towards the robustness of our overall framework in extracting stable associated subspaces from the multimodal neuroimaging data.

Based on group differences, many studies have found the structure and connectivity of default mode network (DMN) being affected in AD and MCI (Greicius et al., 2004; Balachandar et al., 2015; He et al., 2007; Rombouts et al., 2005). (Zhou et al., 2010) also found changes in DMN and saliency networks between AD, MCI, and CN groups. This is also captured in the ASCs obtained for various groups through our method. (Zhang et al., 2012) also analyzed brain regions with the most contribution towards classification into AD/MCI/CN groups, which overlap with the differences in the set of the regions involved in defining ASCs in Figure 5.

More importantly, since sMRI features are known to be more sensitive to Alzheimer’s disease than functional measures, the ASCs associated with the AD group in Figure 5 are primarily defined by structural elements of the multimodal vector rather than the functional ones. Within the structural elements, it can be observed that the contribution could be either negative (diminishing gray matter volume) or positive (increase in gray matter volume) depending on the scores and regions, which could be attributed to atrophy/neuro-degeneration (Pini et al., 2016; Fox and Schott, 2004; Wenk et al., 2003) as well as neuro-inflammation, (Akiyama et al., 2000; Rogers and Shen, 2000; Rogers, 2008) involved in various brain regions in Alzheimer’s disease, respectively.

### 3.6 Performance Comparison

Regression analysis was done by projecting multimodal input features of the randomly selected validation set of each repetition onto the active subspaces from the corresponding repetition. This amounts to compute the transformed feature matrix 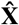 in Equation 5 as input to the regression model. The SVR model with polynomial kernel was trained with 5-fold internal cross-validation on each repetition. Figure 6 shows the Pearson correlation and normalized root mean squared error (NRMSE) between predicted and actual values for all the 7 scores used in the analysis. To compare the predictive performance between the modalities, ASL was done to compute subspaces using only structural, only functional, and multimodal input features, followed by the aforementioned regression analysis on transformed features 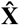 for each modality case. As is clear from Figure 6, multimodal features lead to a significantly better regression performance in almost all cases. Additionally, the performance was comparable to the baseline features without transform, indicating that the predictive information is retained in the active subspaces, with multimodal subspaces retaining the information better than subspaces computed with a single modality. This leads to informed interpretability of the participating components/connections that form each of the active subspace centers.

**Figure 6.**
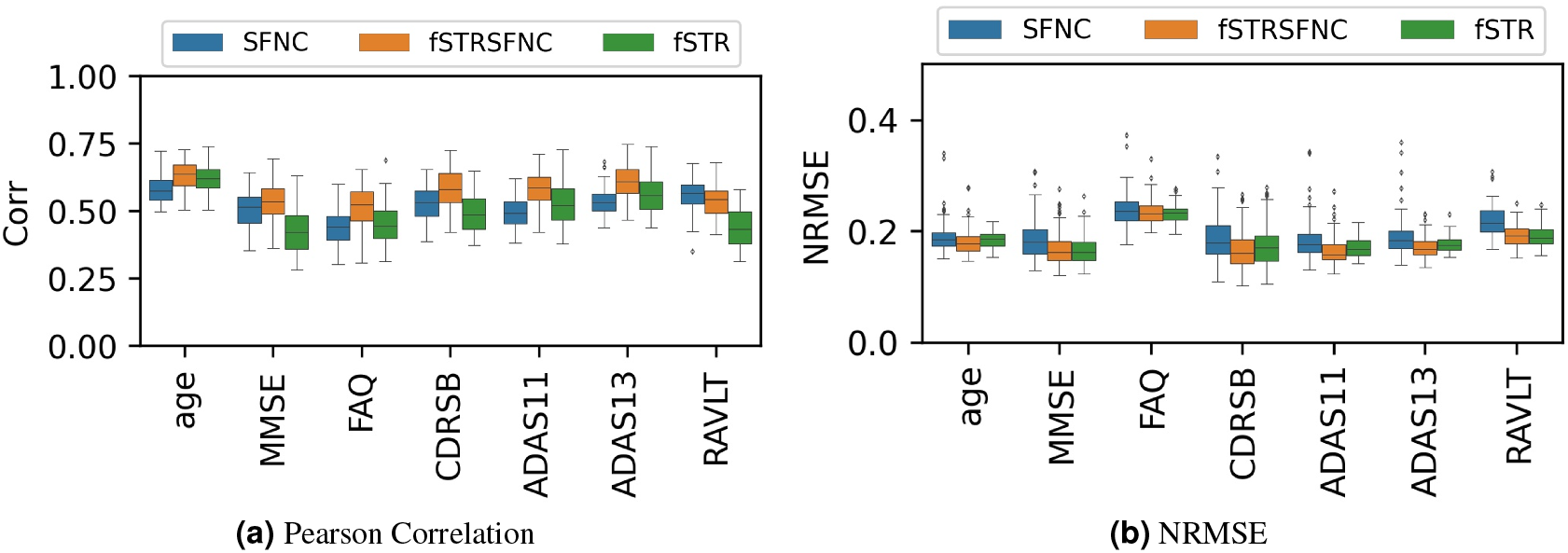
Regression performance of ASL features for 100 repetitions of random sub-sampling with 5-fold cross-validation using SVR with the polynomial kernel. The boxplots show the (a) Pearson correlation and (b) normalized root mean squared error (NRMSE) between the predicted and actual values of the 7 clinical scores for the 100 repetitions. Comparison is shown for active subspaces and projections of features using only static functional connectivity (SFNC), only structural (fSTR), and multimodal input features (fSTRSFNC). It can be noticed that the multimodal features are better at retaining the predictive power of the transformed features than using only structural or only functional features.

## 4 CONCLUSION

Our framework to infer multimodal active subspace patterns can successfully uncover brain subspaces associated with changes in a given cognitive or biological assessment while fusing both structural and functional features. Moreover, since the framework considers the information about the target variable into account, it can retain predictive performance when features projected onto subspaces are used to predict test data. As is clear from the results, various measures and groups of subjects involve different kinds of changes in the underlying subspace patterns in the brain. Thus, rather than looking at the intricate changes at the level of individual features, our framework summarizes the dynamic patterns of modifications in the brain at the level of multimodal subspaces. In the context of Alzheimer’s disease, our framework not only identifies regions that are known to be affected (like hippocampus, entorhinal cortex, DMN), but also summarizes the multiple structural and functional brain sub-systems that characterize the changes in clinically measured outcomes in relation to AD. Such frameworks are essential for the successful development of biomarkers associated with brain disorders. Additionally, this approach can be extended to various deep learning and classification frameworks in the future. To conclude, further research involving the basic and advanced applications of active subspace learning to brain disorders should be conducted.

## DATA/CODE AVAILABILITY STATEMENT

Data used in the preparation of this article were obtained from the Alzheimer’s Disease Neuroimaging Initiative (ADNI) database (adni.loni.usc.edu). The ADNI was launched in 2003 as a public-private partnership led by Principal Investigator Michael W. Weiner, MD. The primary goal of ADNI has been to test whether serial magnetic resonance imaging (MRI), positron emission tomography (PET), other biological markers, and clinical and neuropsychological assessment can be combined to measure the progression of mild cognitive impairment (MCI) and early Alzheimer’s disease (AD).

Data pre-processing was done using SPM12 toolbox (https://www.fil.ion.ucl.ac.uk/spm/software/spm12/), and the scICA components were used from the GIFT toolbox (https://trendscenter.org/software/gift/). The remainder code used for the analysis will be made available upon a simple request to the corresponding author.

## ETHICS AND INFORMED CONSENT

The MRI and fMRI data from human subjects used in this study were from a previously published and publicly available dataset by ADNI. The ethics statement can be found on the ADNI website (adni.loni.usc.edu).

## ACKNOWLEDGEMENTS

Research reported in this publication was supported by the National Institutes of Health (NIH) under award number NIH-RF1AG063153 and by the National Science Foundation (NSF) under award number NSF 2112455.

Data collection and sharing for this project were funded by the Alzheimer’s Disease Neuroimaging Initiative (ADNI) (National Institutes of Health Grant U01 AG024904) and DOD ADNI (Department of Defense award number W81XWH-12-2-0012). ADNI is funded by the National Institute on Aging, the National Institute of Biomedical Imaging and Bioengineering, and through generous contributions from the following: AbbVie, Alzheimer’s Association; Alzheimer’s Drug Discovery Foundation; Araclon Biotech; BioClinica, Inc.; Biogen; Bristol-Myers Squibb Company; CereSpir, Inc.; Cogstate; Eisai Inc.; Elan Pharmaceuticals, Inc.; Eli Lilly and Company; EuroImmun; F. Hoffmann-La Roche Ltd and its affiliated company Genentech, Inc.; Fujirebio; GE Healthcare; IXICO Ltd.; Janssen Alzheimer Immunotherapy Research & Development, LLC.; Johnson & Johnson Pharmaceutical Research & Development LLC.; Lumosity; Lundbeck; Merck & Co., Inc.; Meso Scale Diagnostics, LLC.; NeuroRx Research; Neurotrack Technologies; Novartis Pharmaceuticals Corporation; Pfizer Inc.; Piramal Imaging; Servier; Takeda Pharmaceutical Company; and Transition Therapeutics. The Canadian Institutes of Health Research is providing funds to support ADNI clinical sites in Canada. Private sector contributions are facilitated by the Foundation for the National Institutes of Health (www.fnih.org). The grantee organization is the Northern California Institute for Research and Education, and the study is coordinated by the Alzheimer’s Therapeutic Research Institute at the University of Southern California. ADNI data are disseminated by the Laboratory for Neuro Imaging at the University of Southern California.

## REFERENCES

Abavisani, M. and Patel, V. M. (2018). Deep multimodal subspace clustering networks. IEEE Journal of Selected Topics in Signal Processing, 12(6):1601–1614.

Abrol, A., Bhattarai, M., Fedorov, A., Du, Y., Plis, S., Calhoun, V., Initiative, A. D. N., et al. (2020). Deep residual learning for neuroimaging: an application to predict progression to alzheimer’s disease. Journal of neuroscience methods, 339:108701.

Abrol, A., Fu, Z., Salman, M., Silva, R., Du, Y., Plis, S., and Calhoun, V. (2021). Deep learning encodes robust discriminative neuroimaging representations to outperform standard machine learning. Nature communications, 12(1):1–17.

Adali, T., Akhonda, M., and Calhoun, V. D. (2018). Ica and iva for data fusion: An overview and a new approach based on disjoint subspaces. IEEE sensors letters, 3(1):1–4.

Akiyama, H., Barger, S., Barnum, S., Bradt, B., Bauer, J., Cole, G. M., Cooper, N. R., Eikelenboom, P., Emmerling, M., Fiebich, B. L., et al. (2000). Inflammation and alzheimer’s disease. Neurobiology of aging, 21(3):383–421.

Balachandar, R., John, J., Saini, J., Kumar, K., Joshi, H., Sadanand, S., Aiyappan, S., Sivakumar, P., Loganathan, S., Varghese, M., et al. (2015). A study of structural and functional connectivity in early alzheimer’s disease using rest fmri and diffusion tensor imaging. International journal of geriatric psychiatry, 30(5):497–504.

Bhagwat, N., Pipitone, J., Voineskos, A. N., Chakravarty, M. M., Initiative, A. D. N., et al. (2019). An artificial neural network model for clinical score prediction in alzheimer disease using structural neuroimaging measures. Journal of Psychiatry and Neuroscience, 44(4):246–260.

Braun, U., Schaefer, A., Betzel, R. F., Tost, H. Meyer-Lindenberg, A., and Bassett, D. S. (2018). From maps to multi-dimensional network mechanisms of mental disorders. Neuron, 97(1):14–31.

Bro, R., Acar, E., and Kolda, T. G. (2008). Resolving the sign ambiguity in the singular value decomposition. Journal of Chemometrics: A Journal of the Chemometrics Society, 22(2):135–140.

Bro, R., Leardi, R., and Johnsen, L. G. (2013). Solving the sign indeterminacy for multiway models. Journal of Chemometrics, 27(3-4):70–75.

Calhoun, V. D., Liu, J., and Adalı, T. (2009). A review of group ica for fmri data and ica for joint inference of imaging, genetic, and erp data. Neuroimage, 45(1):S163–S172.

Calhoun, V. D., Pearlson, G. D., and Sui, J. (2021). Data-driven approaches to neuroimaging biomarkers for neurological and psychiatric disorders: emerging approaches and examples. Current Opinion in Neurology, 34(4):469–479.

Calhoun, V. D. and Sui, J. (2016). Multimodal fusion of brain imaging data: a key to finding the missing link (s) in complex mental illness. Biological psychiatry: cognitive neuroscience and neuroimaging, 1(3):230–244.

Chang, Y.-L., Bondi, M., McEvoy, L. Fennema-Notestine, C., Salmon, D., Galasko, D., Hagler, D., Dale, A., Initiative, A. D. N., et al. (2011). Global clinical dementia rating of 0.5 in mci masks variability related to level of function. Neurology, 76(7):652–659.

Constantine, P. G., Dow, E., and Wang, Q. (2014). Active subspace methods in theory and practice: applications to kriging surfaces. SIAM Journal on Scientific Computing, 36(4):A1500–A1524.

Du, Y., Fu, Z., Sui, J., Gao, S., Xing, Y., Lin, D., Salman, M., Abrol, A., Rahaman, M. A., Chen, J., et al. (2020). Neuromark: An automated and adaptive ica based pipeline to identify reproducible fmri markers of brain disorders. NeuroImage: Clinical, 28:102375.

Erhardt, E. B., Rachakonda, S., Bedrick, E. J., Allen, E. A., Adali, T., and Calhoun, V. D. (2011). Comparison of multi-subject ica methods for analysis of fmri data. Human brain mapping, 32(12):2075–2095.

Folstein, M. F., Folstein, S. E., and McHugh, P. R. (1975). “mini-mental state”: a practical method for grading the cognitive state of patients for the clinician. Journal of psychiatric research, 12(3):189–198.

Fornito, A., Zalesky, A., and Breakspear, M. (2015). The connectomics of brain disorders. Nature Reviews Neuroscience, 16(3):159–172.

Fox, N. C. and Schott, J. M. (2004). Imaging cerebral atrophy: normal ageing to alzheimer’s disease. The Lancet, 363(9406):392–394.

Golbabaei, S., Dadashi, A., and Soltanian-Zadeh, H. (2016). Measures of the brain functional network that correlate with alzheimer’s neuropsychological test scores: An fmri and graph analysis study. In 2016 38th Annual International Conference of the IEEE Engineering in Medicine and Biology Society (EMBC), pages 5554–5557. IEEE.

Gomar, J. J., Bobes-Bascaran, M. T., Conejero-Goldberg, C., Davies, P., Goldberg, T. E., Initiative, A. D. N., et al. (2011). Utility of combinations of biomarkers, cognitive markers, and risk factors to predict conversion from mild cognitive impairment to alzheimer disease in patients in the alzheimer’s disease neuroimaging initiative. Archives of general psychiatry, 68(9):961–969.

Greicius, M. D., Srivastava, G., Reiss, A. L., and Menon, V. (2004). Default-mode network activity distinguishes alzheimer’s disease from healthy aging: evidence from functional mri. Proceedings of the National Academy of Sciences, 101(13):4637–4642.

He, Y., Wang, L., Zang, Y., Tian, L., Zhang, X., Li, K., and Jiang, T. (2007). Regional coherence changes in the early stages of alzheimer’s disease: a combined structural and resting-state functional mri study. Neuroimage, 35(2):488–500.

Hughes, C. P., Berg, L., Danziger, W., Coben, L. A., and Martin, R. L. (1982). A new clinical scale for the staging of dementia. The British journal of psychiatry, 140(6):566–572.

Jafri, M. J., Pearlson, G. D., Stevens, M., and Calhoun, V. D. (2008). A method for functional network connectivity among spatially independent resting-state components in schizophrenia. Neuroimage, 39(4):1666–1681.

Klöppel, S., Abdulkadir, A., Jack Jr, C. R., Koutsouleris, N., Mourão-Miranda, J., and Vemuri, P. (2012). Diagnostic neuroimaging across diseases. Neuroimage, 61(2):457–463.

Liu, N., Zong, Y., Zhang, B., Liu, L., Chen, J., Zhao, G., and Zhu, J. (2018). Unsupervised cross-corpus speech emotion recognition using domain-adaptive subspace learning. In 2018 IEEE International Conference on Acoustics, Speech and Signal Processing (ICASSP), pages 5144–5148. IEEE.

Lv, J., Kang, Z., Lu, X., and Xu, Z. (2021). Pseudo-supervised deep subspace clustering. IEEE Transactions on Image Processing, 30:5252–5263.

Ma, H., Li, E.-P., Cangellaris, A. C., and Chen, X. (2020). Support vector regression-based active subspace (svr-as) modeling of high-speed links for fast and accurate sensitivity analysis. IEEE Access, 8:74339–74348.

María Mateos-Pérez, J., Dadar, M., Lacalle-Aurioles, M., Iturria-Medina, Y., Zeighami, Y., and Evans, A. C. (2018). Structural neuroimaging as clinical predictor: a review of machine learning applications. arXiv e-prints, pages arXiv–1806.

Molloy, D. W. and Standish, T. I. (1997). A guide to the standardized mini-mental state examination. International psychogeriatrics, 9(S1):87–94.

Moradi, E., Hallikainen, I., Hänninen, T., Tohka, J., Initiative, A. D. N., et al. (2017). Rey’s auditory verbal learning test scores can be predicted from whole brain mri in alzheimer’s disease. NeuroImage: Clinical, 13:415–427.

Mwangi, B., Tian, T. S., and Soares, J. C. (2014). A review of feature reduction techniques in neuroimaging. Neuroinformatics, 12(2):229–244.

Nielsen, A. N., Barch, D. M., Petersen, S. E., Schlaggar, B. L., and Greene, D. J. (2020). Machine learning with neuroimaging: Evaluating its applications in psychiatry. Biological Psychiatry: Cognitive Neuroscience and Neuroimaging, 5(8):791–798.

Nyberg, L., Boraxbekk, C.-J., Sörman, D. E., Hansson, P., Herlitz, A., Kauppi, K., Ljungberg, J. K., Lövheim, H., Lundquist, A., Adolfsson, A. N., et al. (2020). Biological and environmental predictors of heterogeneity in neurocognitive ageing: evidence from betula and other longitudinal studies. Ageing research reviews, 64:101184.

Perneczky, R., Hartmann, J., Grimmer, T., Drzezga, A., and Kurz, A. (2007). Cerebral metabolic correlates of the clinical dementia rating scale in mild cognitive impairment. Journal of geriatric psychiatry and neurology, 20(2):84–88.

Pini, L., Pievani, M., Bocchetta, M., Altomare, D., Bosco, P., Cavedo, E., Galluzzi, S., Marizzoni, M., and Frisoni, G. B. (2016). Brain atrophy in alzheimer’s disease and aging. Ageing research reviews, 30:25–48.

Razzaghi, P., Razzaghi, P., and Abbasi, K. (2019). Transfer subspace learning via low-rank and discriminative reconstruction matrix. Knowledge-based systems, 163:174–185.

Rogers, J. (2008). The inflammatory response in alzheimer’s disease. Journal of periodontology, 79:1535–1543.

Rogers, J. and Shen, Y. (2000). A perspective on inflammation in alzheimer’s disease. Annals of the New York Academy of Sciences, 924(1):132–135.

Rombouts, S. A., Barkhof, F., Goekoop, R., Stam, C. J., and Scheltens, P. (2005). Altered resting state networks in mild cognitive impairment and mild alzheimer’s disease: an fmri study. Human brain mapping, 26(4):231–239.

Rosen, W. G., Mohs, R. C., and Davis, K. L. (1984). A new rating scale for alzheimer’s disease. The American journal of psychiatry.

Salami, A., Eriksson, J., and Nyberg, L. (2012). Opposing effects of aging on large-scale brain systems for memory encoding and cognitive control. Journal of Neuroscience, 32(31):10749–10757.

Sui, J., Jiang, R., Bustillo, J., and Calhoun, V. (2020). Neuroimaging-based individualized prediction of cognition and behavior for mental disorders and health: methods and promises. Biological psychiatry, 88(11):818–828.

Sun, Z., Chiong, R., and Hu, Z.-p. (2018). An extended dictionary representation approach with deep subspace learning for facial expression recognition. Neurocomputing, 316:1–9.

Turan, C., Zhao, R., Lam, K.-M., and He, X. (2021). Subspace learning for facial expression recognition: an overview and a new perspective. APSIPA Transactions on Signal and Information Processing, 10.

Wan, J., Zhang, Z., Yan, J., Li, T., Rao, B. D., Fang, S., Kim, S., Risacher, S. L., Saykin, A. J., and Shen, L. (2012). Sparse bayesian multi-task learning for predicting cognitive outcomes from neuroimaging measures in alzheimer’s disease. In 2012 IEEE Conference on Computer Vision and Pattern Recognition, pages 940–947. IEEE.

Wang, R., Li, Y., and Darve, E. (2018). On the numerical rank of radial basis function kernels in high dimensions. SIAM Journal on Matrix Analysis and Applications, 39(4):1810–1835.

Wang, Z., Nie, F., Tian, L., Wang, R., and Li, X. (2020). Discriminative feature selection via a structured sparse subspace learning module. In IJCAI, pages 3009–3015.

Wenk, G. L. et al. (2003). Neuropathologic changes in alzheimer’s disease. Journal of Clinical Psychiatry, 64:7–10.

Woo, C.-W., Chang, L. J., Lindquist, M. A., and Wager, T. D. (2017). Building better biomarkers: brain models in translational neuroimaging. Nature neuroscience, 20(3):365–377.

Wycoff, N., Binois, M., and Wild, S. M. (2021). Sequential learning of active subspaces. Journal of Computational and Graphical Statistics, 30(4):1224–1237.

Xu, L., Groth, K. M., Pearlson, G., Schretlen, D. J., and Calhoun, V. D. (2009). Source-based morphometry: The use of independent component analysis to identify gray matter differences with application to schizophrenia. Human brain mapping, 30(3):711–724.

Zhang, D., Shen, D., Initiative, A. D. N., et al. (2012). Multi-modal multi-task learning for joint prediction of multiple regression and classification variables in alzheimer’s disease. NeuroImage, 59(2):895–907.

Zhang, W. and Song, P. (2019). Transfer sparse discriminant subspace learning for cross-corpus speech emotion recognition. IEEE/ACM Transactions on Audio, Speech, and Language Processing, 28:307–318.

Zhou, J., Greicius, M. D., Gennatas, E. D., Growdon, M. E., Jang, J. Y., Rabinovici, G. D., Kramer, J. H., Weiner, M., Miller, B. L., and Seeley, W. W. (2010). Divergent network connectivity changes in behavioural variant frontotemporal dementia and alzheimer’s disease. Brain, 133(5):1352–1367.

