## Supplementary Materials for "SVR-based Multimodal Active Subspace Analysis for the Brain using Neuroimaging Data"

### 1. DATA DEMOGRAPHICS

| | age | MMSE | FAQ | CDRSB | ADAS11 | ADAS13 | RAVLT |
|---|---|---|---|---|---|---|---|
| mean | 73.79 | 27.1 | 4.27 | 1.63 | 10.63 | 16.85 | 35.07 |
| std | 7.0 | 2.65 | 6.2 | 1.75 | 6.25 | 9.37 | 12.4 |

**Table S1.** Mean and standard deviation (std) of all the scores used in the dataset. The seven scores are age, Mini-Mental State Exam (MMSE), Functional Activities Questionnaire (FAQ), Clinical Dementia Rating Scale Sum of Boxes (CDRSB), Alzheimer's Disease Assessment Scale (ADAS11, ADAS13), and Rey's Auditory Verbal Learning Test (RAVLT).

### 2. SVR-ASL WITH RBF AND LINEAR KERNELS

Based on the methodology laid out in sections 2.2 and 2.3 in the main manuscript, it is technically possible to also use the radial basis function (RBF) or the linear kernel for the SVR-ASL process. For the RBF kernel with width $\sigma > 0$, the SVR prediction function and the corresponding gradient vector can be computed for plugging into the steps for the SVR-ASL defined in sections 2.2, 2.3 of the paper as follows:

$$f(\mathbf{u}) = \sum_{1\le i\le n} (\alpha_i^{'} - \alpha_i)\ \exp\left(-\frac{1}{2\sigma^2}||\mathbf{u} - \mathbf{X_i}||^2\right) + b \tag{S1}$$

$$\frac{\partial f(\mathbf{X})}{\partial \mathbf{u}_k} = (\nabla_{\mathbf{u}} f)_k = \sum_{1\le i\le n} (\alpha_i^{'} - \alpha_i)\ \left(-\frac{1}{\sigma^2}(\mathbf{u}_k - \mathbf{X}_{ik})\right) \exp\left(-\frac{1}{2\sigma^2}||\mathbf{u} - \mathbf{X_i}||^2\right) \tag{S2}$$

While the RBF kernel is commonly used in SVR-related applications, in our particular case, the exponential term in Equation S2 becomes very close to zero due to floating point precision issues and high dimensions of the vector involved, leading to almost null gradients. Thus we rejected the RBF kernel as an option for the SVR-ASL analysis.

However, instead of estimating the matrix **C** as sample covariance of data points, one could also attempt to derive a theoretical expression for the covariance matrix **C**, given the gradient expression from the learned SVR-regression function from the data. This avoids sampling the gradient values for data points corresponding to the subjects or augmented synthetic data. The theoretical expression cannot be computed easily for SVR with polynomial or radial kernels but can be computed for SVR with the linear kernel as below:

$$f(\mathbf{u}) = \sum_{1\le i\le n} (\alpha_i^{'} - \alpha_i)\ \mathbf{X}_i^T \mathbf{u} + b \tag{S3}$$

$$\frac{\partial f(\mathbf{u})}{\partial \mathbf{u}_k} = \sum_{1\le i\le n} (\alpha_i^{'} - \alpha_i)\mathbf{X}_{ik} = (\mathbf{\Delta\alpha})^T \mathbf{X}_{*k}\ ,\ \text{ where } \mathbf{X}_{*k} \text{ is the } k^{th} \text{ column of } \mathbf{X} \tag{S4}$$

$$\left[(\nabla_u f)(\nabla_u f)^T\right] = (\mathbf{\Delta\alpha})^T \mathbf{X}\mathbf{X}^T \mathbf{\Delta\alpha} \tag{S5}$$

$$i.e.\ \left[(\nabla_u f)(\nabla_u f)^T\right]_{jk} = \sum_{1\le i\le n} (\alpha_i^{'} - \alpha_i)\mathbf{X}_{ik} \tag{S6}$$

$$\mathbf{C} = \mathbb{E}_{\mathbf{u}}\left[(\nabla_{\mathbf{u}} f)(\nabla_{\mathbf{u}} f)^T\right] = \int_{\mathbf{u}} \left((\mathbf{\Delta\alpha})^T \mathbf{X}\mathbf{X}^T \mathbf{\Delta\alpha}\right)\ p(\mathbf{u})d\mathbf{u} = (\mathbf{\Delta\alpha})^T \mathbf{X}\mathbf{X}^T \mathbf{\Delta\alpha} \tag{S7}$$

Thus, the matrix **C** can be computed directly using coefficients and support vectors of the learned SVR function instead of estimation from existing or generated data points. However, as noticeable in Equation S4, the gradient term in the case of the linear kernel is constant, i.e., independent of the input vector $u$. Thus, for a given dataset, the SVR model with a linear kernel would give a constant gradient at all points, making it unfit for the ASL analysis.

### 3. CLUSTER CENTROIDS AND ACTIVE SUBSPACE CENTERS

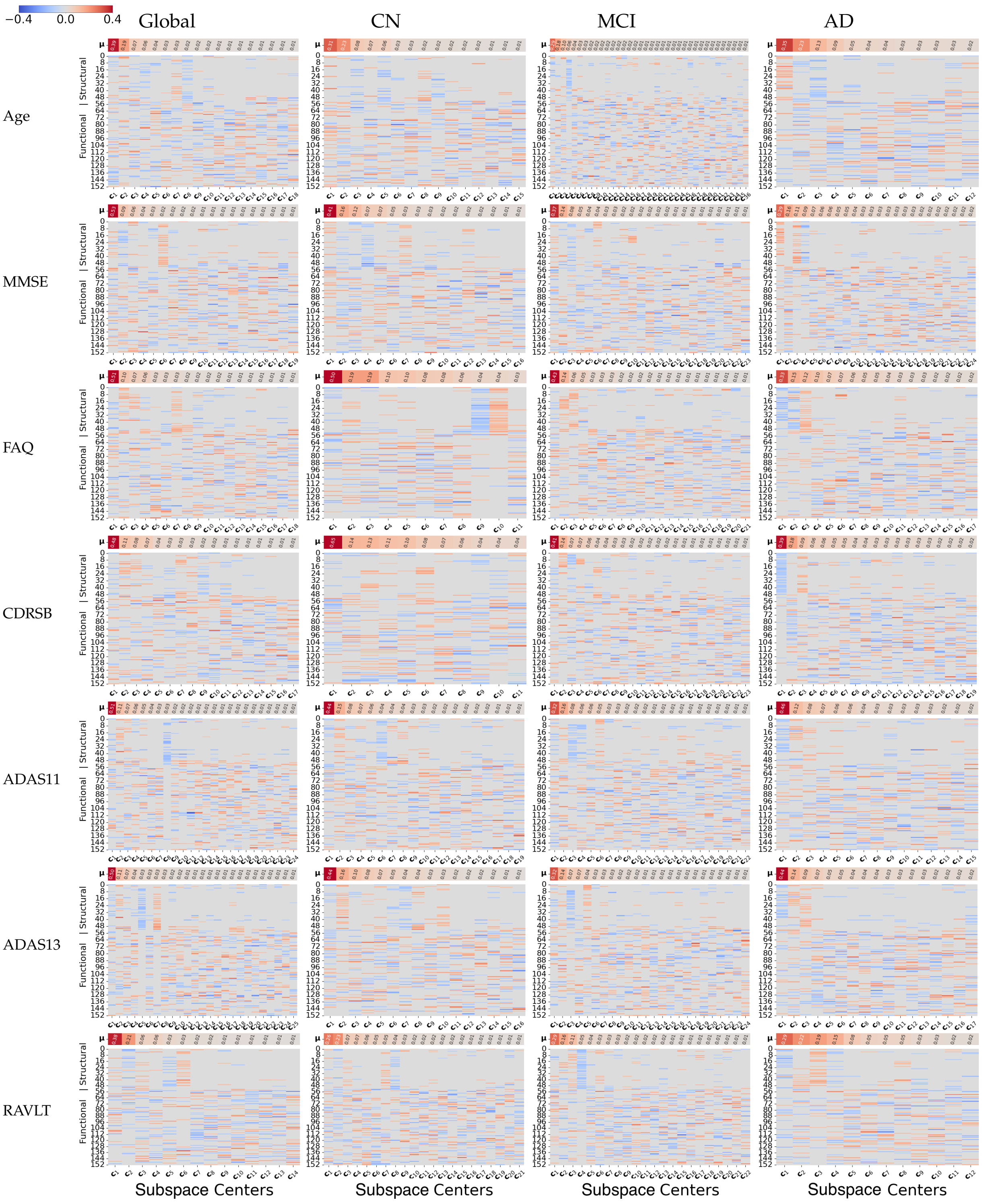


**Fig. S1.** Clustering of active subspace vectors from across 100 repetitions of the ASL analysis. Each column represents a cluster centroid (Active Subspace Center). Top clusters were selected based on mean fractional contribution ($\mu >$ 0.1) and are plotted as connectograms in the main manuscript.
